# A role for worm *cutl-24* in background- and parent-of-origin-dependent ER stress resistance

**DOI:** 10.1101/2021.07.01.450736

**Authors:** Wenke Wang, Anna G. Flury, Andrew T. Rodriguez, Jennifer L. Garrison, Rachel B. Brem

## Abstract

Organisms in the wild can acquire disease- and stress-resistance traits that outstrip the programs endogenous to humans. Finding the molecular basis of such natural resistance characters is a key goal of evolutionary genetics. Standard statistical-genetic methods toward this end can perform poorly in organismal systems that lack high rates of meiotic recombination, like *Caenorhabditis* worms. Here we discovered unique ER stress resistance in a wild Kenyan *C. elegans* isolate, which in inter-strain crosses was passed by hermaphrodite mothers to hybrid offspring. We developed an unbiased version of the reciprocal hemizygosity test, RH-seq, to explore the genetics of this parent-of-origin-dependent phenotype. Among top-scoring gene candidates from a partial-coverage RH-seq screen, we focused on the neuronally-expressed, cuticlin-like gene *cutl-24* for validation. In gene disruption and controlled crossing experiments, we found that *cutl-24* was required in Kenyan hermaphrodite mothers for ER stress tolerance in their inter-strain hybrid offspring; *cutl-24* was also a contributor to the trait in purebred backgrounds. These data establish the Kenyan strain allele of *cutl-24* as a determinant of a natural stress-resistant state, and they set a precedent for the dissection of natural trait diversity in invertebrate animals without the need for a panel of meiotic recombinants.

## Introduction

Understanding mechanisms of trait diversity in organisms from the wild is a central goal of modern genetics. Classical methods toward this end, namely association and linkage mapping, have enabled landmark successes in the field (Flint and Mott 2001). These tools are well-suited to organismal systems in which highly polymorphic, well-mixed recombinant populations exist in the wild or can be generated in the lab. Such features are lacking in the nematode worm *Caenorhabditis*. The laboratory strain of *C. elegans* has been a fixture of the biology literature for decades (Nigon and Félix 2017). Dissecting natural variation in nematodes has proven to be a challenge owing to low rates of polymorphism and meiotic recombination (Hillers and Villeneuve 2003; Andersen *et al*. 2012). Landmark resources have been established to break down these limitations (Li *et al*. 2006; Reddy *et al*. 2009; Rockman and Kruglyak 2009; McGrath *et al*. 2009; Doroszuk *et al*. 2009; Seidel *et al*. 2011; Bendesky *et al*. 2012; Ghosh *et al*. 2012, 2015; Balla *et al*. 2015; Andersen *et al*. 2015; Large *et al*. 2016; Ben-David *et al*. 2017), including recent progress with genome-wide association scans (Webster *et al*. 2019; Na *et al*. 2020; Evans *et al*. 2021). Despite these advances, for many nematode strains and species our access to the full power of natural variation genetics remains limited.

The reciprocal hemizygosity test (Stern 2014) is a strategy to map genotype to phenotype that complements linkage and association approaches. This approach starts with an F1 hybrid formed from the mating of two genetically distinct lines of interest. An induced mutation in the hybrid at a given gene disrupts the allele from each parent in turn, uncovering the other allele in a hemizygous state. The set of hemizygote mutants allow a comparison of phenotypes conferred by the two parents’ alleles, controlled for background and ploidy. Strengths of the method include the ability for trait dissection without a panel of meiotic recombinants, and the potential for genetic insights into traits unique to hybrid backgrounds (heterosis, for example, and imprinting and other parent-of-origin effects; (Comings and MacMurray 2000; Reinhold 2002; Wolf and Wade 2009; Timberlake 2013; Monk *et al*. 2019)). Our group previously established genome-wide reciprocal hemizygosity mapping in yeast (RH-seq; (Weiss *et al*. 2018)), and we reasoned that nematodes could serve as a useful testbed for an extension of the method to multicellular animals.

For a case study using RH-seq, we set out to identify natural stress resistance states in wild worm isolates and to probe their genetics. We chose to focus on ER protein folding quality control, which is essential in eukaryotes for development and stress response, and is a linchpin of diabetes, neurodegenerative disease, and other human disorders (Hebert and Molinari 2007; Roth *et al*. 2010). Surveying *C. elegans* isolates, we characterized a Kenyan strain with an ER stress resistance phenotype, and we discovered a robust parent-of-origin effect in hybrids derived from this strain. To enable the search for determinants of this trait variation, we developed an implementation of RH-seq for the worm which achieved unbiased, partial coverage of the genome. From the results we focused on one gene hit, *cutl-24*, for validation of its impact on ER stress response.

## Results

### Tunicamycin resistance in wild *C. elegans* and its parent-of-origin dependence

Tunicamycin, a N-glycosylation inhibitor and ER stress inducer, causes developmental delay in nematodes (Richardson *et al*. 2011; Shen *et al*. 2001). To identify *C. elegans* strains resistant to this defect, we collected eggs from each of a panel of wild isolates and the laboratory strain N2, exposed them to a toxic concentration of tunicamycin, and tabulated the number of successfully developed adults after 96 hours at 20°C. Among these strains, the predominant phenotype was of marked sensitivity, in which most eggs exposed to tunicamycin failed to reach adulthood, and indeed never reached the larval stage of development (Figure 1). All isolates we assayed fit this description except one: ED3077, originally isolated from a park in Nairobi, Kenya, had a rate of development in the presence of the drug outstripping that of the rest of the panel by >2-fold (Figure 1). We earmarked this tunicamycin resistance phenotype in ED3077 as a compelling target for genetic dissection, and we chose to use N2 as a representative of the tunicamycin-sensitive state.

**Figure 1.**
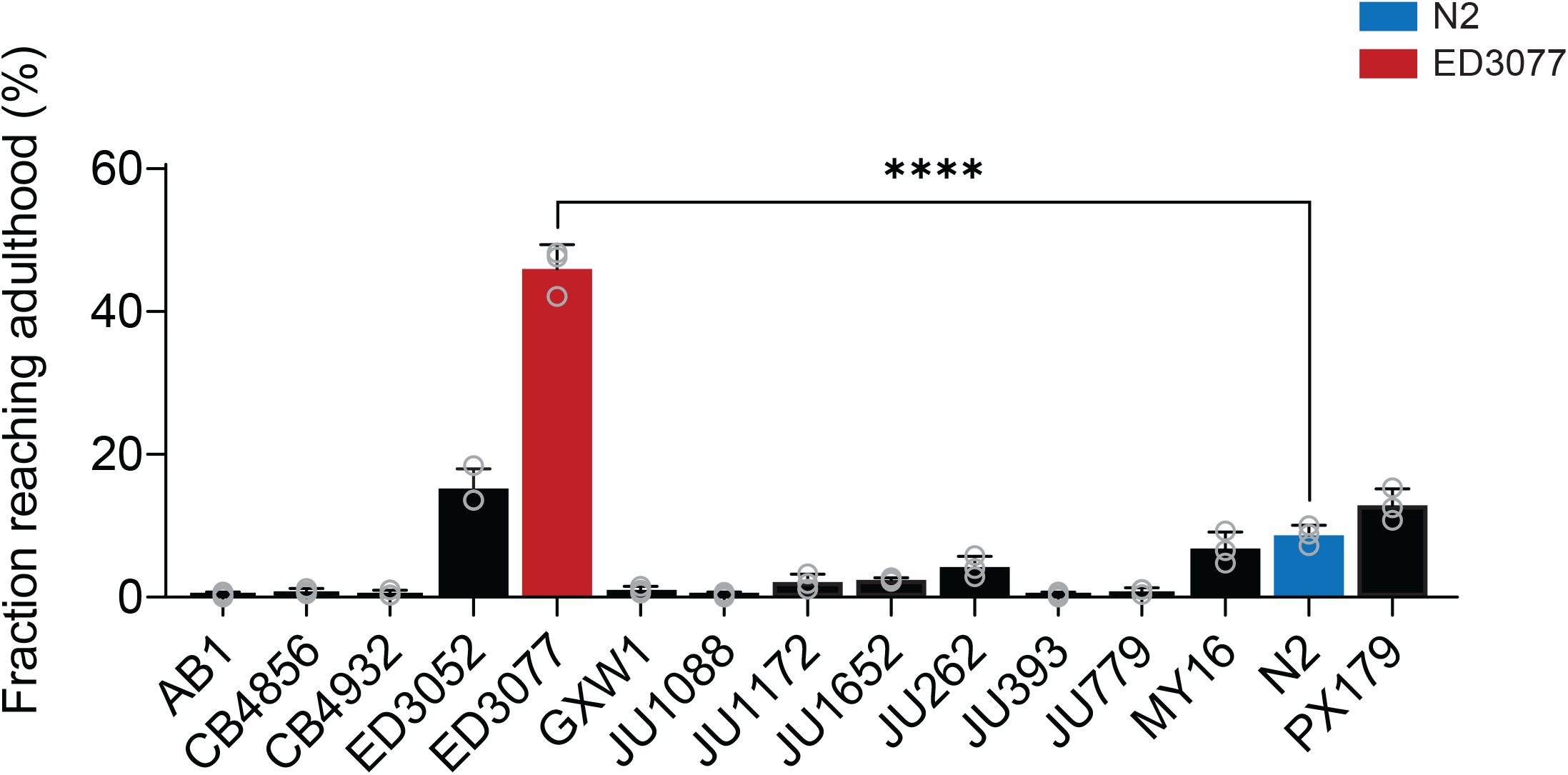
Tunicamycin resistance phenotypes of wild *C. elegans* isolates. The *y*-axis reports the proportion of eggs from the indicated wild isolate that developed to adulthood in the presence of tunicamycin. For a given column, each dot represents results from one replicate population, and the bar height reports the mean. ****, unpaired two-tailed *t*-test *p* < 0.0001.

To begin to investigate the genetics of tunicamycin resistance in ED3077, we mated this strain to N2, and subjected the resulting F1 hybrid eggs to our tunicamycin development assay. Results made clear that the hybrid phenotype depended on the direction of the cross (Figure 2). ED3077 hermaphrodites, mated to N2 males, yielded eggs that developed in tunicamycin conditions at a rate approaching that of the ED3077 purebred (Figure 2, third column). By contrast, hybrids with N2 as the hermaphrodite parent had almost no ability to develop in the presence of tunicamycin (Figure 2, fourth column). Thus, the ED3077 phenotype was partially dominant over that of N2, but only when ED3077 was the hermaphrodite in the cross. We hypothesized that the mechanism of this maternal effect could bear on the genetics of the trait difference between purebred ED3077 and N2. In what follows, we describe our experiments to probe both facets of the system.

**Figure 2.**
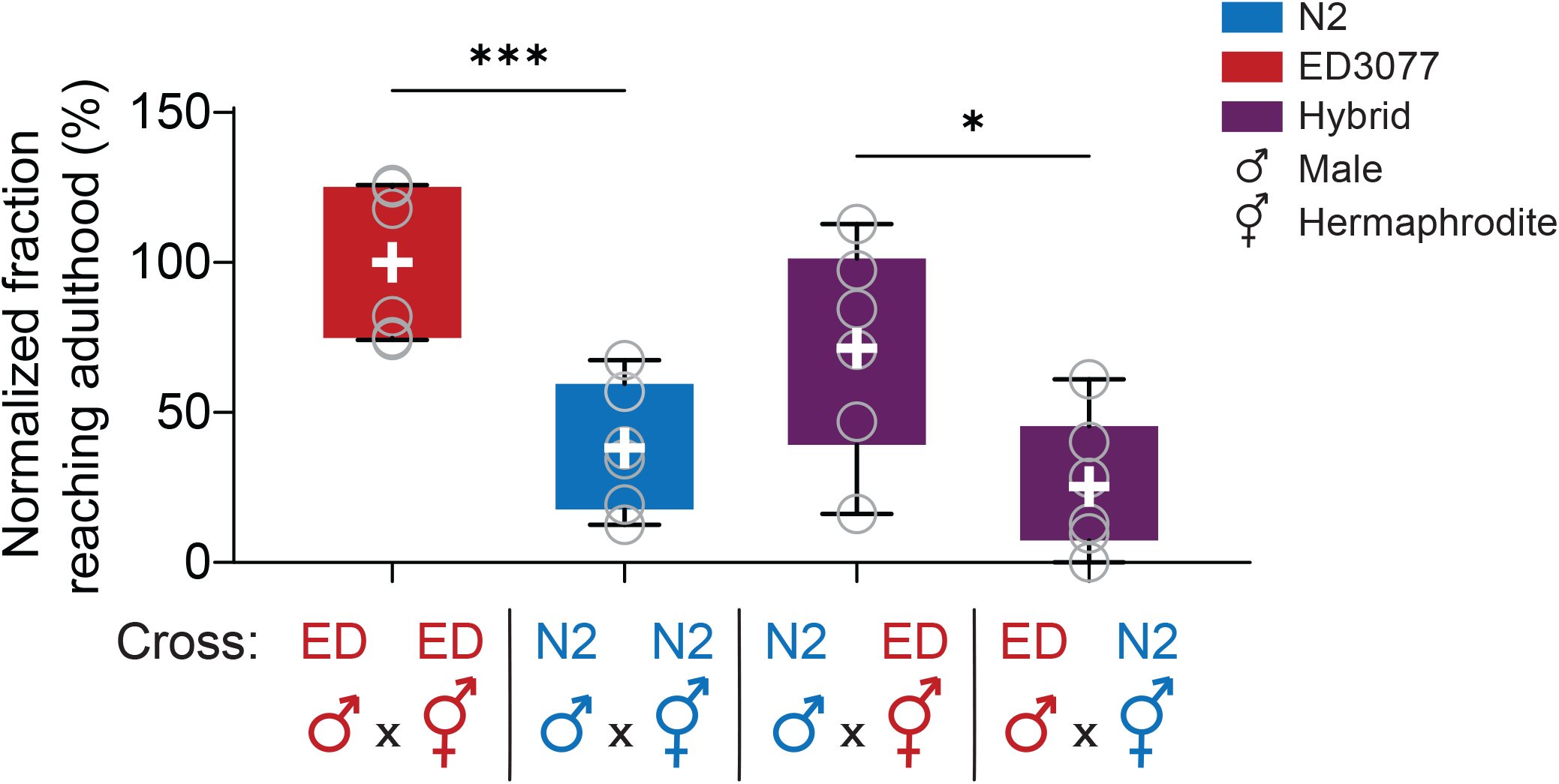
Tunicamycin resistance of inter-strain hybrids depends on the parent of origin. The *y*-axis reports the proportion of eggs from the indicated cross that developed to adulthood in the presence of tunicamycin, normalized to the analogous quantity from wild-type ED3077. For a given column, each dot represents results from one replicate population; the white cross reports the mean; box and whiskers report the interquartile range and the 10-90 percentile range, respectively, of the replicate measurement distribution. ED, ED3077. *, unpaired two-tailed *t*-test *p* < 0.05; ***, *p* < 0.001.

### RH-seq reveals effects of variation in *cutl-24* in an inter-strain hybrid

We sought to investigate the genetic basis of the parent-of-origin-dependent tunicamycin resistance trait in the ED3077 x N2 system using RH-seq. The method requires large cohorts of hybrids harboring disrupting mutations in one of the two copies of each gene in turn, and we chose the Mos1 transposon system (Bessereau *et al*. 2001; Williams *et al*. 2005; Duverger *et al*. 2007) for this purpose. We set up a workflow to generate transposon mutants in ED3077 or N2 and mate each to the wild-type of the other respective parent, yielding hemizygote eggs (Figure 3). In each case we used hermaphrodites as the transposon mutant parent and males as the wild-type, in light of the importance of the hermaphrodite genotype in hybrids (Figure 2). We incubated hemizygote eggs in tunicamycin or untreated control conditions and then collected animals that reached adulthood for DNA isolation and sequencing.

**Figure 3.**
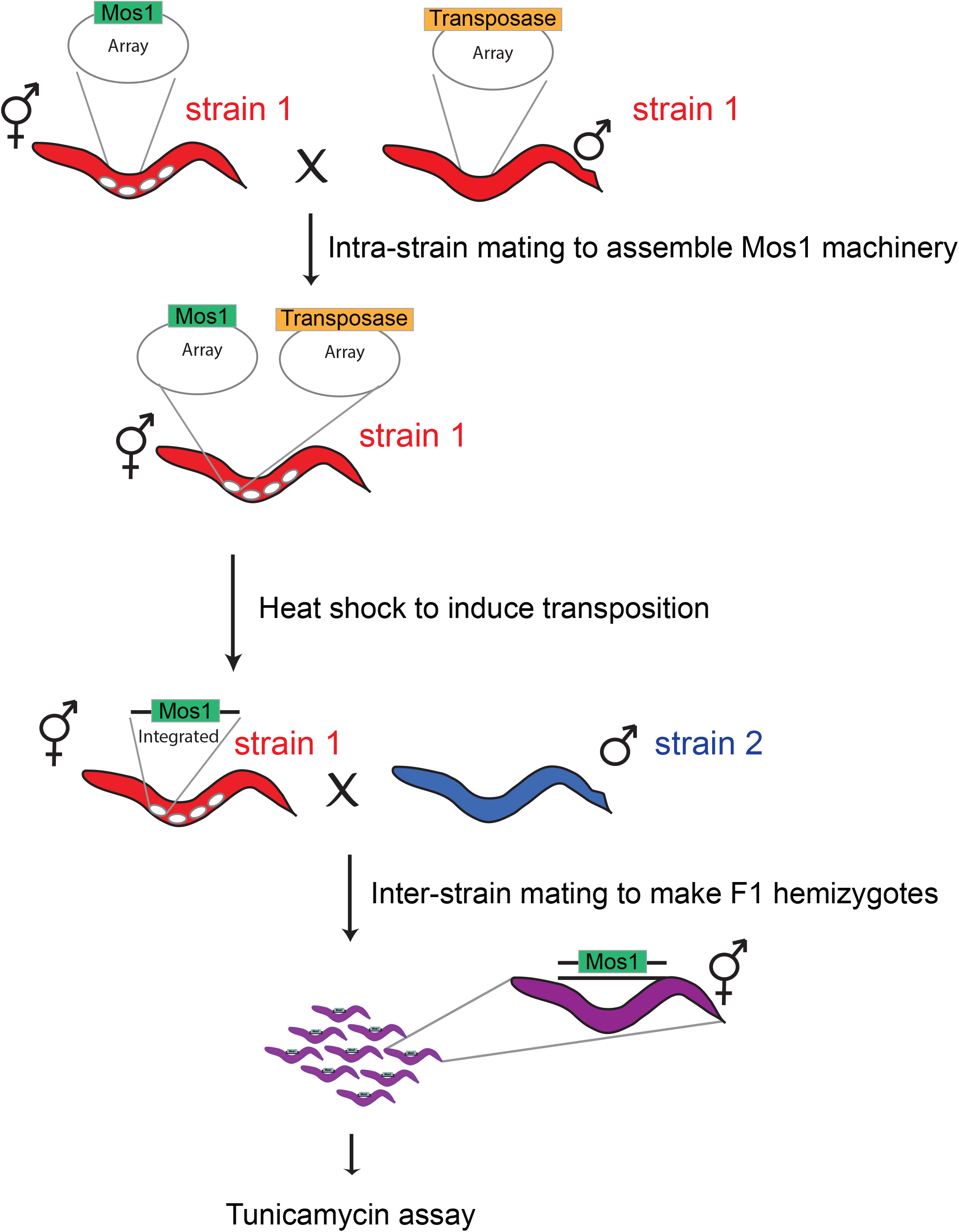
Making hemizygote mutants for RH-seq. RH-seq requires hemizygote hybrids (purple) from crosses between mutants of one background (red) and wild-types of another (blue). Top: arrays (circles) harboring the Mos1 transposon (green) and heat-shock-inducible transposase enzyme gene (orange) in the red background come together into one strain. Center: after heat shock, a transposon copy integrates into the genome (straight black line) of an egg of the red background, which is fertilized by a wild-type male of the blue background. Bottom: the resulting F1 hybrids are hemizygous throughout the soma and are used as input into a sequencing-based tunicamycin resistance assay.

We expected that deploying RH-seq at moderate scale could serve as a proof of concept for the method and help discover candidate determinants of tunicamycin resistance in the ED3077 x N2 hybrid. To this end, we generated and detected a total of 56,979 hemizygote genotypes from DNA of animals reaching adulthood in tunicamycin and in control conditions, after quality-control filtering of sequencing data (Supplementary Tables 2 and 3).

Focusing on the 2,721 highest-coverage genes, for each we assessed the difference in normalized sequencing representation among tunicamycin-treated adults between two cohorts of hemizygotes—those with the ED3077 allele uncovered and those with the N2 allele uncovered. In this test, no results reached significance after correction for multiple testing (Supplementary Figure 1 and Supplementary Table 4). Nonetheless, we reasoned that the top-scoring loci represented our most compelling candidate determinants of tunicamycin response. Manual inspection of the top ten RH-seq hits revealed that for each of the top three, *WBGene00023395/Y82E9BL.9, WBGene00015173/trpp-11*, and *WBGene00021396/cutl-24*, RH-seq data supported the inference of a pro-resistance function for the ED3077 allele (Supplementary Figure 2). That is, at each such locus, genotypes with the ED3077 allele disrupted in the hybrid, leaving the N2 uncovered and functional, were on average at low abundance in sequencing of tunicamycin-treated adults relative to controls; genotypes with the ED3077 allele intact and the N2 allele disrupted were abundant on average in these samples. We considered genes with effects of this direction in RH-seq to hold the most promise in helping explain the tunicamycin resistance of purebred ED3077. Among them, we noted that agreement in RH-seq abundance measures between independent transposon inserts was strongest for *cutl-24*, encoding a largely uncharacterized protein containing a cuticlin-like domain (Figure 4). On this basis, we had the highest confidence in *cutl-24* as a potential determinant of tunicamycin resistance, and we chose it as a target for experiments to validate the role of the gene, and its inter-strain variation, on the trait.

**Figure 4.**
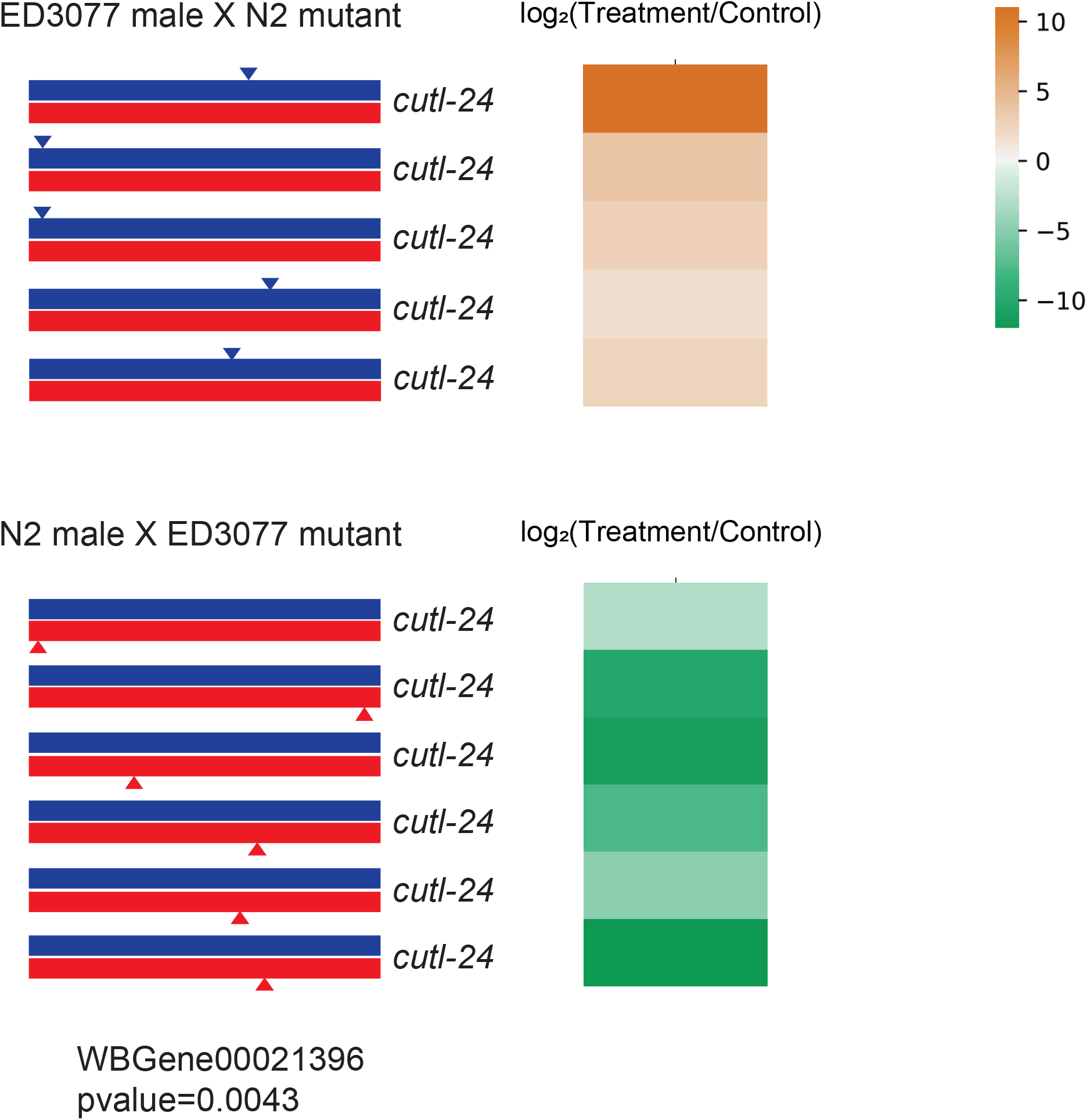
RH-seq reveals *cutl-24* as a candidate gene at which inter-strain variation contributes to tunicamycin resistance. In a given panel, in each row the left-hand cartoon represents the region of *cutl-24* in the hybrid genome (blue line, ED3077 chromosome; red line, N2 chromosome), and the triangle denotes the position of insertion of a Mos1 transposon as detected by transposon sequencing. The right-hand cell reports the log_2_ of the abundance of the respective mutant, detected by sequencing, after development in tunicamycin, relative to the analogous quantity from development in untreated control conditions. The *p*-value reports the result of a two-tailed Mann-Whitney statistical test for a difference in the abundance after tunicamycin selection, relative to the abundance in an untreated control, of hemizygotes harboring transposon insertions in the two parents’ orthologs. Supplementary Table 4 reports Mos1 insertion positions and raw quantitation data, with the Benjamini-Hochberg method used to correct for multiple testing. Top, hemizygotes harboring transposon insertions in the N2 allele; bottom, hemizygotes harboring transposon insertions in the ED3077 allele.

### *cutl-24* as a determinant of tunicamycin resistance in hybrids and purebreds

To investigate the impact of *cutl-24* on tunicamycin response in the ED3077 x N2 system, we first focused on the hybrid background, where this gene had been a top scorer in the RH-seq screen (Figure 4). We used Cas9 to generate stable lines of purebred N2 and ED3077 harboring a nonsense mutation in *cutl-24* (Supplementary Table 5). We mated each mutant as the hermaphrodite to the wild-type male of the respective other strain, as in our RH-seq framework (Figures 3-4), and we subjected the progeny to high-sensitivity development assays in the presence of tunicamycin. The results showed that disrupting *cutl-24* in ED3077 hermaphrodites, followed by crossing to wild-type male N2, compromised tunicamycin resistance of the hybrid by 28.5% (Figure 5a; compare second and first columns), consistent with the trend we had observed in RH-seq (Figure 4). Mutating *cutl-24* in N2 hermaphrodites, followed by mating with wild-type ED3077 males, had no such effect (Figure 5a; compare fourth and third columns). These data establish *cutl-24* as a driver of the unique tunicamycin resistance phenotype of the hybrid progeny of ED3077 hermaphrodites, serving as a validation of the RH-seq approach.

**Figure 5.**
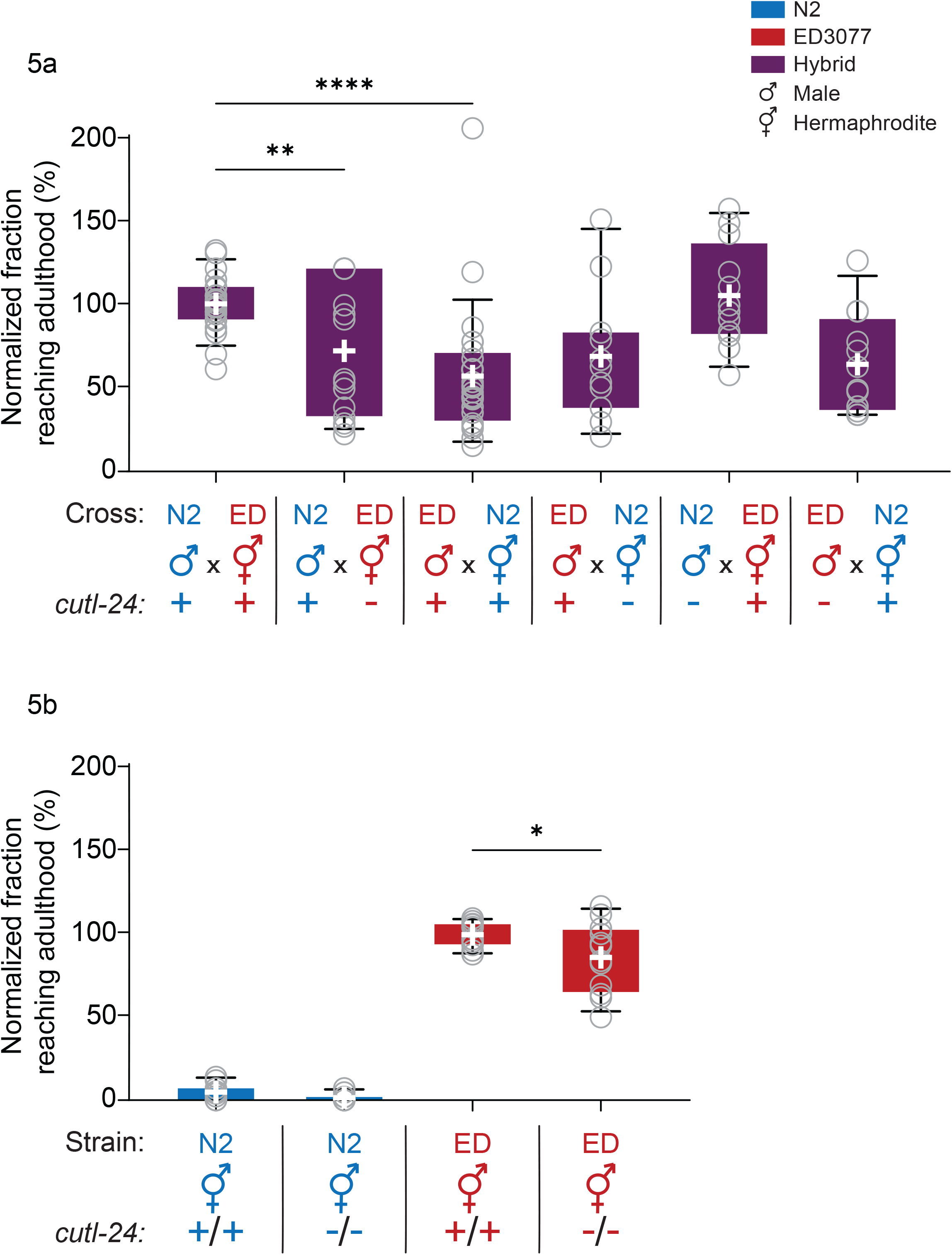
*cutl-24* in ED3077 mothers is required for tunicamycin resistance in their inter-strain hybrid progeny. (a) The *y*-axis reports tunicamycin resistance measurements in F1 hybrid animals from crosses between wild-type or *cutl-24* mutant ED3077 and N2 as indicated, normalized per experiment with respect to the mean value from the wild-type ED3077 hermaphrodite x N2 male F1. (b) Data are as in (a) except that strains were purebred *cutl-24* mutants or wild-type controls, and measurements were normalized to the mean value from wild-type ED3077. For a given column, each dot represents results from one replicate population; the white cross reports the mean; box and whiskers report the interquartile range and the 10-90 percentile range, respectively, of the replicate measurement distribution. ED, ED3077. *, unpaired two-tailed *t*-test *p* < 0.05; **, *p* < 0.01; ****, *p* < 0.0001.

We also examined the importance of *cutl-24* in the male parent of inter-strain hybrids. *cutl-24* mutation in ED3077 males had no detectable effect on tunicamycin resistance of their hybrid progeny from crosses with N2 hermaphrodites (Figure 5a; compare sixth and third columns); the same was true when N2 mutants were the male parent of a cross with ED3077 hermaphrodites (Figure 5a; compare fifth and first columns). This result echoed our finding from wild-type crosses that hermaphrodite genotype matters most for tunicamycin resistance in hybrids (Figure 2). Together, our genetic analyses establish *cutl-24* as a determinant of the parent-of-origin-dependent tunicamycin resistance in ED3077 x N2 hybrids, with the ED3077 allele in the hermaphrodite sufficient to explain the hybrid resistance phenotype entirely.

We hypothesized that *cutl-24* could also contribute to tunicamycin resistance in a purebred context. In purebred ED3077, we observed a 13.9% drop in tunicamycin resistance in the *cutl-24* mutant relative to wild-type (Figure 5b, red); the trend persisted, though not significantly so, in the N2 background, where survival in tunicamycin was almost nil to start with (reduced by 66.7% upon *cutl-24* mutation; Figure 5b, blue). These relatively modest effects of *cutl-24* mutation in purebreds contrasted with the stronger dependence of the resistance phenotype on *cutl-24* in hybrids (Figure 5a), pointing to the hybrid as a sensitized background in which *cutl-24* dependence is amplified and drives appreciable parent-of-origin effects. In purebreds, we infer that *cutl-24* acts as a modifier of tunicamycin resistance, likely one of many determinants of a complex genetic architecture.

### Evidence for *cutl-24* as a neuronal factor

To help formulate models for *cutl-24* activity and function, we characterized the localization of its gene product with an expression analysis approach. In whole-worm developmental timecourses (Dillman *et al*. 2015; Hashimshony *et al*. 2015), *cutl-24* expression was high in embryos and lower but appreciable in larvae and adults (Figure 6). Lineage-traced single-cell transcriptomes through development (Packer *et al*. 2019) detected *cutl-24* in glia of the inner labial sensilla, and, at lower levels, in many other neurons (Figure 7). Likewise, in adult hermaphrodites, RNA tagging for a marker of outer labial sensilla neurons and PVD nociceptors revealed significant *cutl-24* expression (Smith *et al*. 2010). These results establish the presence of the *cutl-24* gene product in a range of neuronal cells and raise the possibility that *cutl-24* may exert its effect on development and stress resistance in this context.

**Figure 6.**
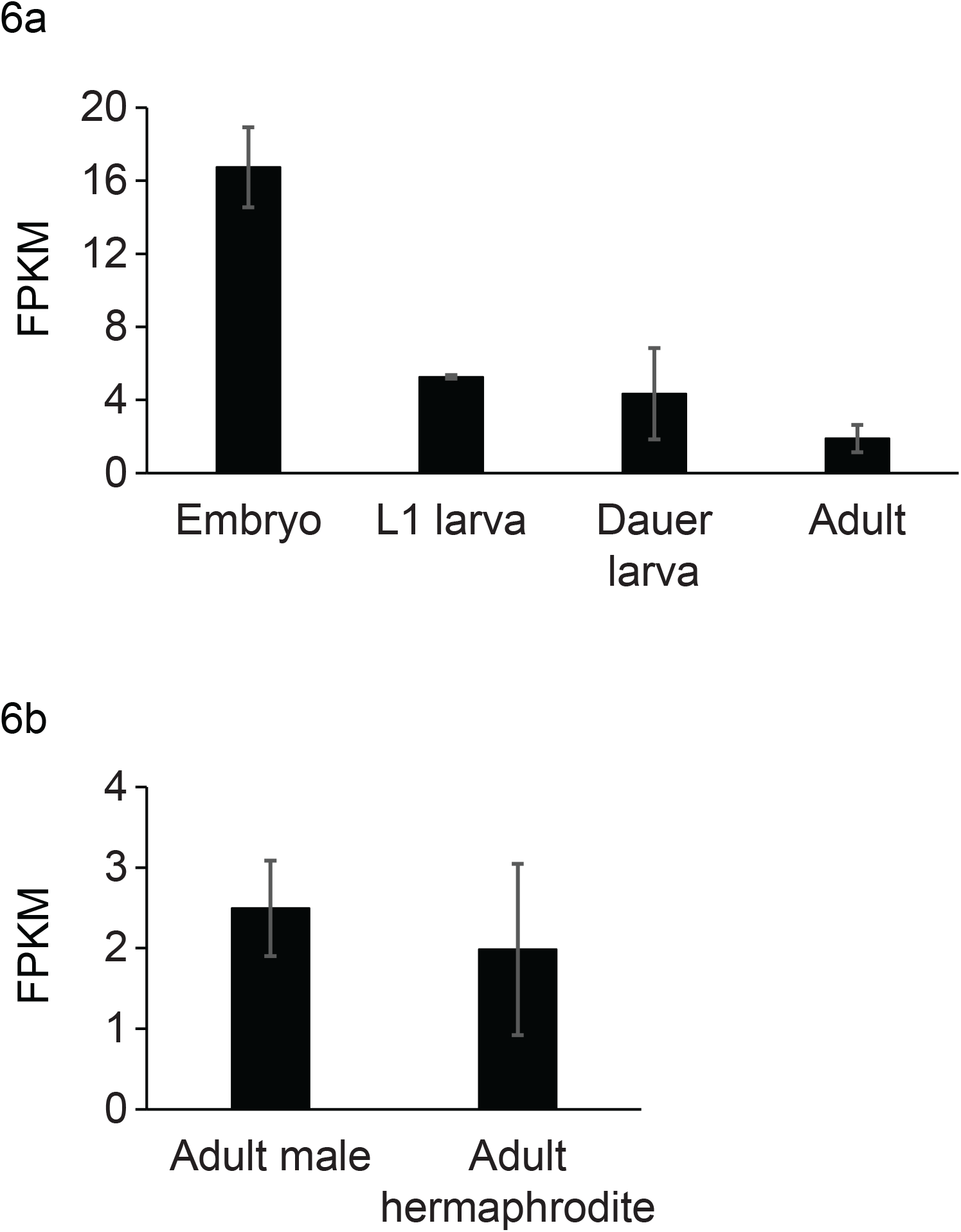
*cutl-24* expression across *C. elegans* developmental stages. In a given panel, each bar reports average expression (FPKM, Fragments Per Kilobase of transcript per Million mapped reads) of *cutl-24* in wild-type animals of the N2 background from (Dillman *et al*. 2015; Hashimshony *et al*. 2015) in the indicated developmental stage (a) or sex of adult animals (b). Error bars report standard deviation (*n* = 2).

**Figure 7.**
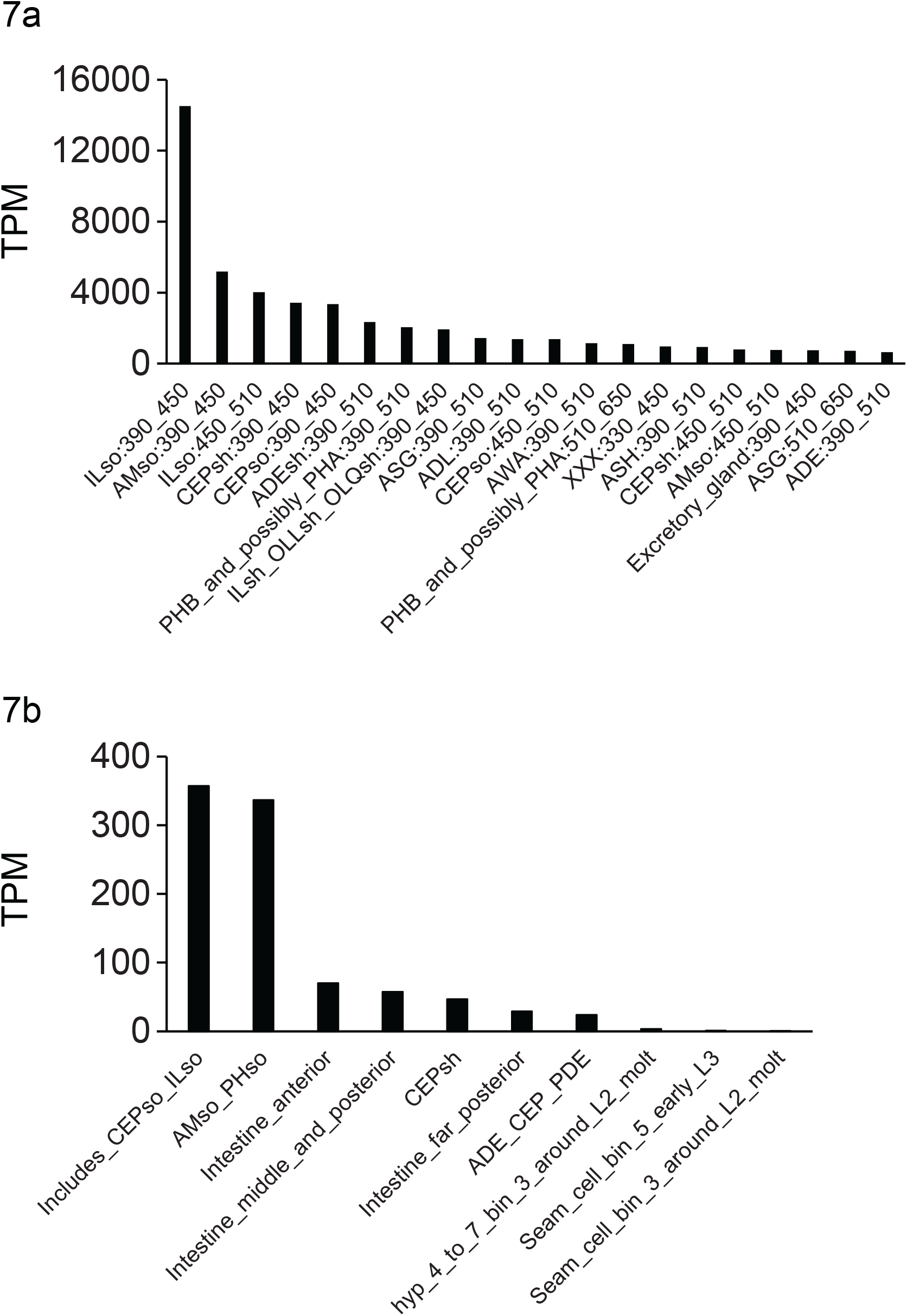
*cutl-24* is expressed in neurons. In a given panel, each bar reports *cutl-24* expression (TPM, Transcripts Per Million kilobases) in the indicated cell type from single cell transcriptomes (Packer *et al*. 2019), in embryos at the indicated developmental time in minutes (a) or in L2 larvae (b). The highest expression was seen in inner labial socket (ILso), cephalic socket (CEPso), and phasmid socket (PHso) cells; for other abbreviations in labels, see Methods.

## Discussion

The search for genes underlying natural trait variation has been a mandate for geneticists since Mendel. Wild nematodes have great potential as a study system for ecological genetics (Frézal and Félix 2015; Cook *et al*. 2017). In these animals, landmark studies have identified variants with Mendelian (de Bono and Bargmann 1998; Reddy *et al*. 2009; McGrath *et al*. 2009; Bendesky *et al*. 2012; Ghosh *et al*. 2012, 2015; Balla *et al*. 2015; Large *et al*. 2016) and maternal- and paternal-effect (Ben-David *et al*. 2017; Ewe *et al*. 2020; Seidel *et al*. 2011) modes of action that underlie phenotypes of interest. Many of these successes have derived from model advanced intercross mapping populations in *C. elegans* (Li *et al*. 2006; Rockman and Kruglyak 2009; Doroszuk *et al*. 2009; Andersen *et al*. 2015). In non-model nematode strains and species, statistical genetics continues to pose challenges, owing to population structure among wild isolates (Webster *et al*. 2019) and low meiotic recombination rates in lab crosses (Hillers and Villeneuve 2003).

In this work, we have established strain differences and parent-of-origin effects in worm ER stress resistance. And we have used a new tool, RH-seq, alongside classical genetics to identify *cutl-24* as a determinant of the variation. By discovering maternal-effect and stress-tolerance functions of this uncharacterized gene, our study underscores the utility of the natural variation approach, and RH-seq in particular, in nematodes.

### The RH-seq approach in *C. elegans*

RH-seq shares with several other worm screening methods (Frézal *et al*. 2018; Burga *et al*. 2019; Webster *et al*. 2019) the ability to streamline phenotyping assays by bulk sequencing of naturally varying strains. The chief distinction of RH-seq is its ability to map genotype to phenotype without recombinant progeny from crosses, in the lab or in the wild (Weiss *et al*. 2018; Weiss and Brem 2019). Here the technique has enabled partial-coverage genetic dissection in non-model *C. elegans* ED3077, in the absence of a panel of recombinant inbred lines. More broadly, the utility of RH-seq for any worm strains or species will be a function of mutagenesis throughput and phenotyping signal-to-noise ratio, which will govern coverage and statistical power. Our mapping of *cutl-24* in a screen of limited depth and power serves as a first proof of concept of the method.

### Maternal effects of strain variation and *cutl-24* function

Starting from the discovery of a unique tunicamycin resistance phenotype in the Kenyan *C. elegans* isolate ED3077, we found that ED3077 x N2 hybrid embryos develop best in tunicamycin when they arise from ED3077 hermaphrodite parents rather than ED3077 males. These hybrid progeny rely on parent *cutl-24* more than do ED3077 purebreds, under tunicamycin treatment. Plausibly, hybrids could act as a sensitized background for perturbations in *cutl-24* owing to stresses from genome incompatibility, as has been documented in other *C. elegans* crosses (Dolgin *et al*. 2007; Snoek *et al*. 2014). If so, hermaphrodite *cutl-24* could help maintain homeostasis under this burden, when compounded by tunicamycin.

Our work also leaves open the mechanism by which *cutl-24* in hermaphrodites affects developing progeny. Given its expression in multiple neuronal types, *cutl-24* could participate in the pathway linking neuronal serotonin in mothers with stress-protective transcription in their progeny (Das *et al*. 2020). Alternatively, acting in labial sensilla in particular, *cutl-24* could follow the precedent from other maternal-effect genes that directly modulate egg-laying and egg structure in the worm (Daniele *et al*. 2020; Min *et al*. 2020; Baugh and Hu 2020). Or *cutl-24* transcripts could be expressed in the germline and maternally deposited in the embryo for activity early in development, as is the case for other well-characterized maternal-effect factors (Robertson and Lin 2015). As one potential clue to the mechanism of *cutl-24* genetics, we noted that there was no identifiable DNA sequence variation between ED3077 and N2 in genome resequencing data or Sanger sequencing, in or near the gene. It is thus tempting to speculate that a heritable epigenetic polymorphism, dependent on parent sex and background, could drive the differences in phenotypic impact we have seen between the strains’ alleles at *cutl-24* and contribute to the overall parent-of-origin effects we have noted in ED3077 x N2 crosses. Such a mechanism at *cutl-24* would dovetail with the classic literature describing paramutation in maize (Pilu 2015) and the more modern mapping of epialleles to phenotype in a number of systems (Quadrana *et al*. 2014; Bertozzi *et al*. 2021; Pignatta *et al*. 2018).

Under any of these scenarios, ED3077 hermaphrodites, reared under standard conditions in our assay design, would lay eggs better equipped to handle environmental and/or genetic challenges, in part mediated by *cutl-24*. Consistent with this notion, in a previous study of wild *C. elegans*, the ED3077 strain was distinguished by a marked ability for L1 larvae to develop successfully after starvation (Webster *et al*. 2019). The emerging picture is one of a unique, broad stress-resistant phenotype in ED3077. Further work will establish its complete genetic architecture and the evolutionary forces that drove its appearance in the wild.

## Acknowledgements

This work was funded by NIH R01 GM120430 to R.B.B. and J.L.G., by NIA T32 AG052374 to A.T.R., and by a Glenn Foundation fellowship to W.W. The authors thank Jean-Louis Bessereau and Jonathan Ewbank for their generosity with unpublished resources, and Christian Frøkjær-Jensen, Marie Delattre, and Marie-Anne Félix for discussions.

## Materials and methods

### Worm strains and maintenance

Strains and plasmids used in this work are listed in Supplementary Table 1. For maintenance and crossing, worms were maintained on nematode growth media (NGM) plates seeded with *Escherichia coli* OP50. Tunicamycin resistance assays and RH-seq screening (see below) used 5X OP50 and 1.5X agar to inhibit burrowing. All incubations were at 20°C except as described. Wild-type strains of *C. elegans* AB1 (Adelaide, Australia), CB4856 (Hawaii, USA), CB4932 (Taunton, Great Britain), ED3052 (Ceres, South Africa), ED3077 (Nairobi, Kenya), GXW1 (Wuhan, China), JU1088 (Kakegawa, Japan), JU1172 (Concepcion, Chile), JU1652 (Montevideo, Uruguay), JU262 (Indre, France), JU393 (Hermanville, France), JU779 (Lisbon, Portugal), MY16 (Munster, Germany), N2 (Bristol, Great Britain) and PX179 (Eugene, USA) were used in this study, all from the *Caenorhabditis* Genetics Center.

### High-sensitivity tunicamycin resistance assays

Tunicamycin resistance assays were performed as described (Henis-Korenblit et al. 2010) with modifications as follows. For Figure 1, 2-3 gravid adults were placed onto NGM plates, which were prepared with 5 μg/mL tunicamycin in DMSO (InSolution, Sigma) or DMSO at the analogous percentage by volume for control plates prior to plate pouring, and allowed to lay eggs for 12 hours, after which the adults were removed from the plates. The number of eggs was counted (typically 50-200) and compared with the number of animals that reached the adult stage within 96 hours at 20°C. On control plates, ~100% of eggs hatched for a given strain. For each bar of Figure 2, 5-10 L4 hermaphrodites of one strain were incubated with 10-20 males of another strain for 24 hours to allow mating. The hermaphrodites were then transferred onto plates containing 5 μg/mL tunicamycin (InSolution, Sigma) as above and allowed to lay eggs for 12 hours. The proportion of eggs from the indicated cross that developed to adulthood in the presence of tunicamycin was measured as above and was normalized to the analogous quantity from wild-type ED3077. Experiments for Figure 5 were as Figure 2 with the following changes: 10 μg/mL tunicamycin (InSolution, Sigma) was used, and the number of eggs that reached the adult stage, on the egg-lay tunicamycin plate or on a separate tunicamycin plate to which ~100 eggs were transferred, was counted after 66-68 hours at 25°C, to match the conditions used in high-throughput RH-seq phenotyping (see below).

### RH-seq in *C. elegans* and high-throughput tunicamycin resistance screening

#### RH-seq parental strain construction

To enable the heat-induced Mos1 transposon system (Bessereau *et al*. 2001; Williams *et al*. 2005; Duverger *et al*. 2007) in the ED3077 and N2 backgrounds, we proceeded as follows.

For mutagenesis in N2, we used IG358 (*oxEx229* [Mos1 transposon +P*myo-2::GFP*]), an N2 transgenic which carries multiple copies of the Mos1 transposon and pharynx-specific green fluorescent protein (GFP) expression, and IG444 (*frEx113* [(pJL44 =P*hsp-16.48::Mos1* transposase) + P*col-12::* DsRed]), an N2 transgenic which carries the coding sequence of the Mos1 transposase under the control of a heat-shock promoter and an epidermis-specific DsRed reporter.

For the corresponding strains in the ED3077 background, we amplified the Mos1 transposon used in IG358 (from a plasmid kindly provided by Christian Frøkjaer-Jensen) and cloned it into the pSM plasmid backbone to create pWW1. We then injected pWW1 at 25 ng/μL, pCFJ421 (P*myo-2::gfp::h2b::tbb-2utr*) at 10 ng/μL, and pSM at 65 ng/μL to yield JAZ418. We injected pJL44 (P*hsp16.48::Mos1* transposase*::glh-2utŕ*) at 50 ng/μL, pCFJ90 (P*myo-2::mCherry::unc-54utr*) at 2 ng/μL, and pSM at 48ng/μL to create JAZ419.

Separately, for wild-type marked strains to be used as males in the generation of hemizygotes (see below), wild-type ED3077 and N2 were injected at 5 ng/μL with pCFJ104 (P*myo-3::mCherry::unc-54utr*) and pCFJ90 (P*myo-2::mCherry::unc-54utr*), after which the respective transgene was integrated into the worm genome by UV/TMP treatment (60 μg/mL TMP and 30 mJ of UV exposure at 365 nm) and the worms were outcrossed 3 times, generating JAZ420 and JAZ421, respectively.

#### RH-seq hemizygote construction and phenotyping

Our RH-seq workflow proceeded in 12 rounds in N2. For each round of hemizygotes bearing mutations in the N2 alleles of genes, we first made Mos1-ready N2 animals by picking 100-200 L4 hermaphrodites of IG358 and 200 L4 males of IG444 onto one NGM plate. After 12 hours at 25°C, the hermaphrodites and males were transferred to a new NGM plate for egg-lay. 100-200 F1 progeny exhibiting GFP and mCherry marker expression (*i.e*., containing both transposon and transposase arrays) were picked at the L4 stage.

In these Mos1-ready N2 animals, we induced Mos1 transposition at the young adult stage by heat-shock treatment essentially as described (34°C for 1□h, 20°C for 1 h and then 34°C for 1□h) (Boulin and Bessereau 2007). We collected the putatively mutant eggs and selfed them for two generations to ensure appreciable numbers of clones of a given mutant genotype. We collected 20,000-50,000 F3 progeny and sorted in a Union Biometrica COPAS Biosort for those that had lost the transposon array and transposase array (*i.e*., they lacked GFP and mCherry expression) to prevent continued transposition for the rest of the experiment.

We transferred 10,000-20,000 of these putatively mutant N2 hermaphrodites to NGM plates with 5,000-10,000 JAZ420 males (marked, non-mutated ED3077) and allowed mating to occur for 1.5 days at 15°C. The mated hermaphrodites were lysed, and ~30,000 eggs were transferred to one plate (5-10 plates per round) containing 10 μg/mL tunicamycin, followed by incubation for development for 66-68 hours at 25°C.

Afterward, 3,000-12,000 surviving worms were collected and sorted, with only hermaphrodite adults (assessed based on size) and hybrid progeny of the marked ED3077 parent (assessed based on mCherry marker expression) retained for DNA isolation and sequencing (see below). 11 rounds of hemizygotes bearing mutations in the ED3077 alleles of genes proceeded as above, except that we mated JAZ418 with JAZ419 to make Mos1-ready ED3077, and after mutagenesis, we mated the resulting putatively mutant hermaphrodites in the ED3077 background to JAZ421 males (marked, non-mutated N2) to yield hemizygotes.

### Genomic DNA extraction from *C. elegans*

Animals from hemizygote pools that had been collected after control or tunicamycin treatment from a given RH-seq round were collected and snap-frozen in liquid nitrogen. Genomic DNA (gDNA) from the worm pellets was extracted using the DNeasy Blood & Tissue Kit (Qiagen) and a *C. elegans* gDNA extraction protocol from the Kaganovich lab (University Medical Center Göttingen, http://www.kaganovichlab.com/celegans.html). The purified gDNA was quantified using a Nanodrop.

### Transposon insertion sequencing library construction

Illumina transposon sequencing libraries were constructed using the FS DNA Library Kit (NEB) as follows. By following the protocol, gDNA was enzymatically digested and ligated with E-adapters (a duplex whose upper strand is longer than the lower). The E-adapter upper arm was GGGCGTAGATTACCGTCCGCGACTCGTACTGTGGCGCGCC*T (*T indicates T overhang), and the E-adapter lower arm was /Phos/GGCGCGCCACAGTACTTGACTGAGCTTTA/3ddC/. Sequences containing the 3’ junction of each inserted Mos1 transposon to the genome were amplified using a Mos1 transposon primer (5’- AATGATACGGCGACCACCGAGATCTACACTCTTTCCCTACACGACGCTCTTCCGATCTNNN NNNXXXXXXGATTTAAAAAAAACGACATTTCATAC-3’ N, random sequence; XXXXXX, barcode indices; GATTTAAAAAAAACGACATTTCATAC, Mos1 transposon sequences) and a primer homologous to a region of the adapter (CAAGCAGAAGACGGCATACGAGATACATCGGGGGCGTAGATTACCGTCCGCGACTC). The thermocycler protocol was as follows: 98°C for 30 seconds, (98°C for 10 seconds, 65°C for 1 minute and 15 seconds) x 24, 65°C for 10 minutes, 4°C hold. Single-end sequencing of 100 bp was then done on a NovaSeq machine at the Vincent J. Coates Genomics Sequencing Lab at the University of California, Berkeley. In total, we sequenced 22 libraries from pools of hemizygotes bearing mutations in the ED alleles of genes (representing one tunicamycin-treated and one untreated pool from each of 11 rounds), and 24 libraries from pools bearing mutations in the N2 alleles of genes (representing one tunicamycin-treated and one untreated pool from each of 12 rounds), generating ~4-20 million reads per library (Supplementary Table 2).

### RH-seq read-mapping

For a given library of reads reporting transposon insertion sequencing from hemizygotes after tunicamycin or control treatment for RH-seq, we first removed adapter sequences from the end of the reads using fastx_clipper from the fastx toolkit. Then, using fastx_trimmer from the fastx toolkit, we removed the first 6 bp of every read, so every read would now begin with the barcode index sequence. We then demultiplexed the barcode indexes using fastx_barcode_splitter from the fastx toolkit. Using a custom python script, we retained for analysis only reads containing the last 28 bp of the Mos1 transposon, allowing for no mismatches. For this subset of reads containing the transposon, we excised the sequence immediately flanking the end of the transposon sequence and mapped the remainder of the read (containing the stretch of genome adjoining the transposon insertion) to the respective genome using BLAT (https://genome.ucsc.edu/cgi-bin/hgBlat) with minimum sequence identity = 100 and tile size= 12.

For pools of hemizygotes bearing transposon insertions in the N2 alleles of genes, we mapped to the N2 genome from BioProject PRJNA13758, release WS271. For hemizygotes with insertions in the ED3077 alleles, we mapped to a ED3077 reference genome made as follows. We downloaded raw genome sequencing reads for ED3077 from https://www.elegansvariation.org/ (Cook *et al*. 2017). These reads were aligned to genome assembly WS271 of the reference sequence of the *C. elegans* N2 strain using bowtie2 (http://bowtie-bio.sourceforge.net/bowtie2/manual.shtml) with default parameters. Samtools, bcftools, and bgzip were used to call SNPs, retaining those with a quality score of >20 and combined depth of >5 and <71. We then generated a pseudogenome by replacing the reference N2 allele with that of ED3077 at each SNP using bcftools.

We eliminated reads whose genomic sequence portion was shorter than 30 bp and/or mapped to more than one location in the genome. For analysis of a given sequencing library, we inferred that reads that mapped to genomic positions within 100 bp of each other originated from the same transposon mutant clone and assigned the sum of their counts to the position which had the most mapped reads. Then, for a given insertion in a given library, we considered the number of reads mapped to that position, n_insert_, as the relative proportional abundance of the mutant in the worm pellet whose gDNA was sequenced. In order to compare abundances across samples and libraries, we divided n_insert_ by the total number of reads in the sample to get the normalized abundance.

### RH-seq data analysis

We used N2 genome annotations from BioProject PRJNA13758, release WS271, to determine whether an inferred transposon insertion position was genic or non-coding, and we retained only those insertions which were genic. For a given insertion observed in sequencing from a given RH-seq round (comprising a pool of hemizygotes split into tunicamycin- and control-treated subpools), we tabulated the log_2_ ratio of the normalized abundance in the control sample divided by the normalized abundance in the tunicamycin-treatment sample, which we call the tunicamycin effect on the respective transposon mutant as detected by RH-seq. In formulating this ratio, we assigned a pseudocount of 1 for each case of an insertion with zero reads in either the tunicamycin-treated or control sample of its respective round. For insertions detected in multiple rounds, we averaged the tunicamycin effect across all rounds before proceeding. For a given gene, we compiled all transposon insertion mutants in the gene as observed across all RH-seq rounds, and we used a two-sample Mann-Whitney test to compare the tunicamycin effects between two sets of mutants: those in the N2 allele of the respective gene and those in the ED3077 allele. We corrected for multiple testing using the Benjamini-Hochberg method. One gene from the raw screen data, WBGene00013255, was eliminated from further consideration owing to ambiguity in its annotation across WormBase genome releases.

### Generating and phenotyping Cas9 mutants of *cutl-24*

CRISPR-mediated genome editing was performed as described (Friedland *et al*. 2013) with modifications as follows. We first cloned sgRNA sequences into the pUC57 vector backbone using EcoRI and HindIII restriction enzymes to yield *PU6::cutl-24* sgRNA plasmids, pWW2 and pWW3 for the N2 background (sgRNA sequences GGACAAAGACACACAAACGT and GCAGGCTCCAATAAAGCCGG, respectively) and pWW4 and pWW5 for the ED3077 background (sgRNA sequences GCTGAGATTCGAGgtaagtg, and GACAAAACTTCAAAGATAAC, respectively; uppercase, exonic sequence; lowercase, intronic sequence; see also Supplementary Table 1). Wild-type N2 and ED3077 strains as day 1 adult hermaphrodites were injected with pDD162 (P*eft-3::Cas9::tbb-2utr*) at 50 ng/μL, pCFJ90 (P*myo-2::mCherry*) at 5 ng/μL, and pWW2 and pWW3 or pWW4 and pWW5 respectively, each at 100 ng/μL. Surviving worms were separated, and F1 mCherry-positive animals were collected; their progeny, representing the F2 generation, were genotyped for the *cutl-24* gene. Mutant genotypes (*cutl-24*(*jlg1*) and *cutl-24*(*jlg2*[ED3077]), respectively) are reported in Supplementary Table 5. The resulting N2 and ED3077 *cutl-24* mutants were subsequently crossed with the marked N2 and ED3077 strains JAZ421 and JAZ420 (see *RH-seq parental strain construction* above) to generate JAZ422 and JAZ423, respectively.

For inter-strain crosses in Figure 5a, we mated marked wild-type (JAZ421 for N2 background, JAZ420 for ED3077 background) or *cutl-24* mutant (JAZ422 for N2 background, JAZ423 for ED3077 background) males to marked wild-type or *cutl-24* mutant hermaphrodites of the other respective strain background (ED3077 or N2). We transferred eggs from these crosses to tunicamycin or control plates and carried out development and scoring as above, except that any adult hermaphrodite progeny observed under a fluorescence microscope to lack the respective male marker (P*myo-3::mCherry* from an ED3077 background parent or P*myo-2::mCherry* from an N2 background parent) were censored from the plate as non-hybrid, and the count of eggs (used as the denominator when calculating the final proportion of developed animals) was reduced correspondingly. All adult male progeny were considered to be hybrids due to the link between mating and the generation of male progeny. Purebred worms in Figure 5b were generated by selfing and assayed as above.

### *cutl-24* expression analysis

For Figure 6, the *cutl-24* expression pattern was obtained from (Dillman *et al*. 2015; Hashimshony *et al*. 2015). For Figure 7, the *cutl-24* expression pattern in single cell resolution was obtained from (Packer *et al*. 2019). Cell type labels follow the format CellType:StartTime_EndTime, where times represent development following fertilization: ILso, Inner labial socket; AMso, Amphid socket; CEPsh, Cephalic sheath; CEPso, Cephalic socket; ADEsh, Anterior dereid sheath; PHB_and_possibly_PHA, PHB/PHA neuron; ILsh_OLLsh_OLQsh, Inner/outer labial sheath; XXX, Glia and excretory cells; ASG, ADL, AWA, ASH and ADE, neurons of other types; AMso_PHso, Amphid and phasmid socket; ADE_CEP_PDE, neurons.

### Availability of data and materials

RH-seq data have been deposited in the Sequence Read Archive (SRA), submission ID SUB9564728. Custom Python and R scripts used for RH-seq data analysis are available at https://github.com/annagflury/RHseq-scripts.

## Ethics declarations

The authors declare that they have no competing interests.

## Supplementary figure captions

**Supplementary Figure 1. Genomic distribution of significance measures from RH-seq mapping of tunicamycin resistance.** Each point reports results from a reciprocal hemizygosity test for the impact, at one gene, of variation between ED3077 and N2 on development of their F1 hybrid in the presence of tunicamycin. The *x*-axis reports genome position of the respective gene, and the *y*-axis reports the negative log_10_ of the *p*-value from a Mann-Whitney test comparing two sets of sequencing-based measurements of hybrid strain abundance after development in tunicamycin: those from hybrid hemizygotes bearing a disruption in the ED3077 allele of the gene, uncovering the N2 allele, and those from hemizygotes bearing a disruption in the N2 allele (see Methods). Results for the focal gene of this study, *cutl-24*, are denoted in red.

**Supplementary Figure 2. Top-scoring loci from RH-seq mapping of tunicamycin resistance.** Data are as in Figure 4 of the main text except that each panel reports results from the top 10 most significant genes from reciprocal hemizygosity tests for the impact of variation between ED3077 and N2 on development of their F1 hybrid in the presence of tunicamycin.

## Supplementary table captions

**Supplementary Table 1. Strains and plasmids used in this work.**

**Supplementary Table 2. RH-seq library sizes.** Each row reports sequencing results from hemizygotes in the N2 x ED3077 diploid hybrid background, made from transposon mutants of the indicated strain mated to the wild-type of the other.

**Supplementary Table 3. Abundance of hemizygote mutants in tunicamycin RH-seq.** Abundances of transposon-mutant Mos1 insertion site in the F1 hybrid of N2 or ED3077 allele from RH-seq. Each row reports results of sequencing one transposon insertion in the N2 x ED3077 diploid hybrid after selection of the transposon mutant pool, reflecting the abundance in the pool of the respective hemizygote clone harboring the insertion. Gene, chromosome, and position report the fine-scale position of the insertion. Allele, the strain parent’s homolog in which the transposon insertion lay. Control_count and treatment_count report read counts of the transposon insertion sequenced after selection of the mutant pool in the indicated condition, normalized for library size. Transposon insertions not detected in any replicate of the indicated selection were assigned an abundance of 1. Round reports the batch of Mos1 mutants in which the respective mutant was detected (1-12 for mutagenesis in the N2 parent, 1-11 for mutagenesis in the ED3077 parent).

**Supplementary Table 4. Effects of inter-strain variation in tunicamycin RH-seq.** Each row reports the results of reciprocal hemizygote tests of tunicamycin resistance of hemizygote transposon mutants at the indicated gene in the N2 x ED3077 diploid hybrid. N2_control_count, N2_treatment_count, ED3077_control_count, and ED3077_treatment_count report normalized abundances of a hemizygote harboring a transposon insertion in the indicated parent’s homolog after culture in the indicated condition, as a mean across transposon mutants, from all biological replicates. The last two columns report results of a two-tailed Mann-Whitney statistical test for a difference in the abundance after tunicamycin selection, relative to the abundance in an untreated control, of hemizygotes harboring transposon insertions in the two parents’ homologs. The Benjamini-Hochberg method was used to correct for multiple testing.

**Supplementary Table 5. *cutl-24* mutants.** Each pair of rows reports the context of a Cas9-induced mutation (red) in the *cutl-24* gene of the indicated *C. elegans* strain. Uppercase, exonic sequence; lowercase, intronic sequence.

