## Supplementary figures and images for "A role for worm *cutl-24* in background- and parent-of-origin-dependent ER stress resistance"

### Figure S1

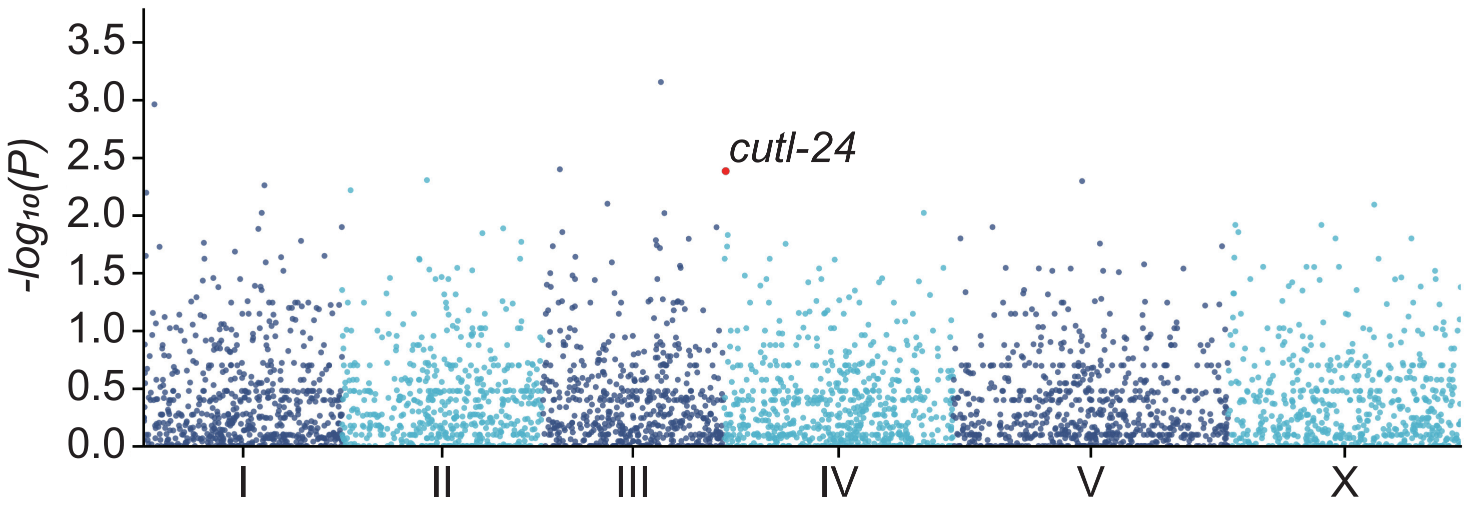


Figure S1

### Figure S2

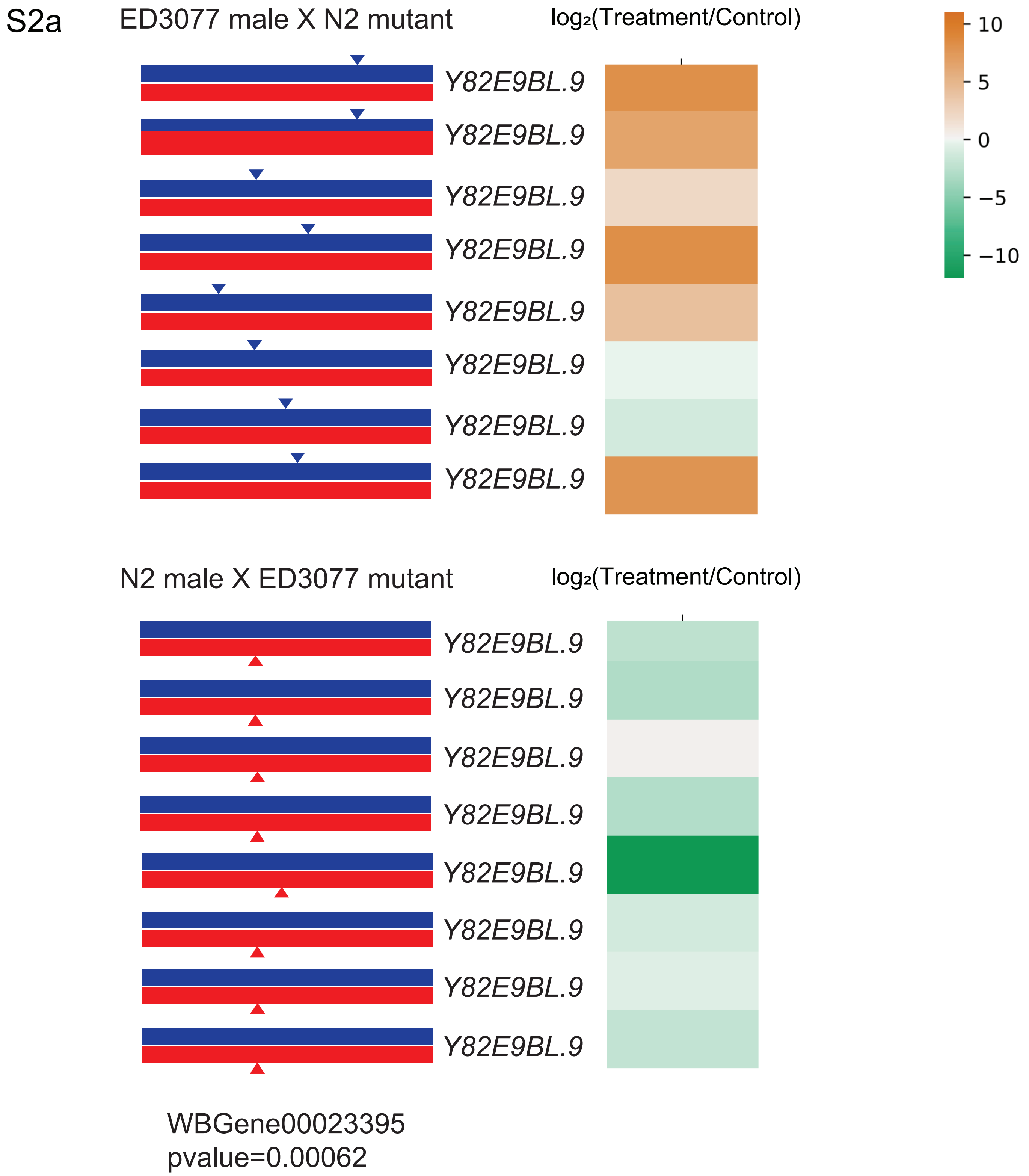

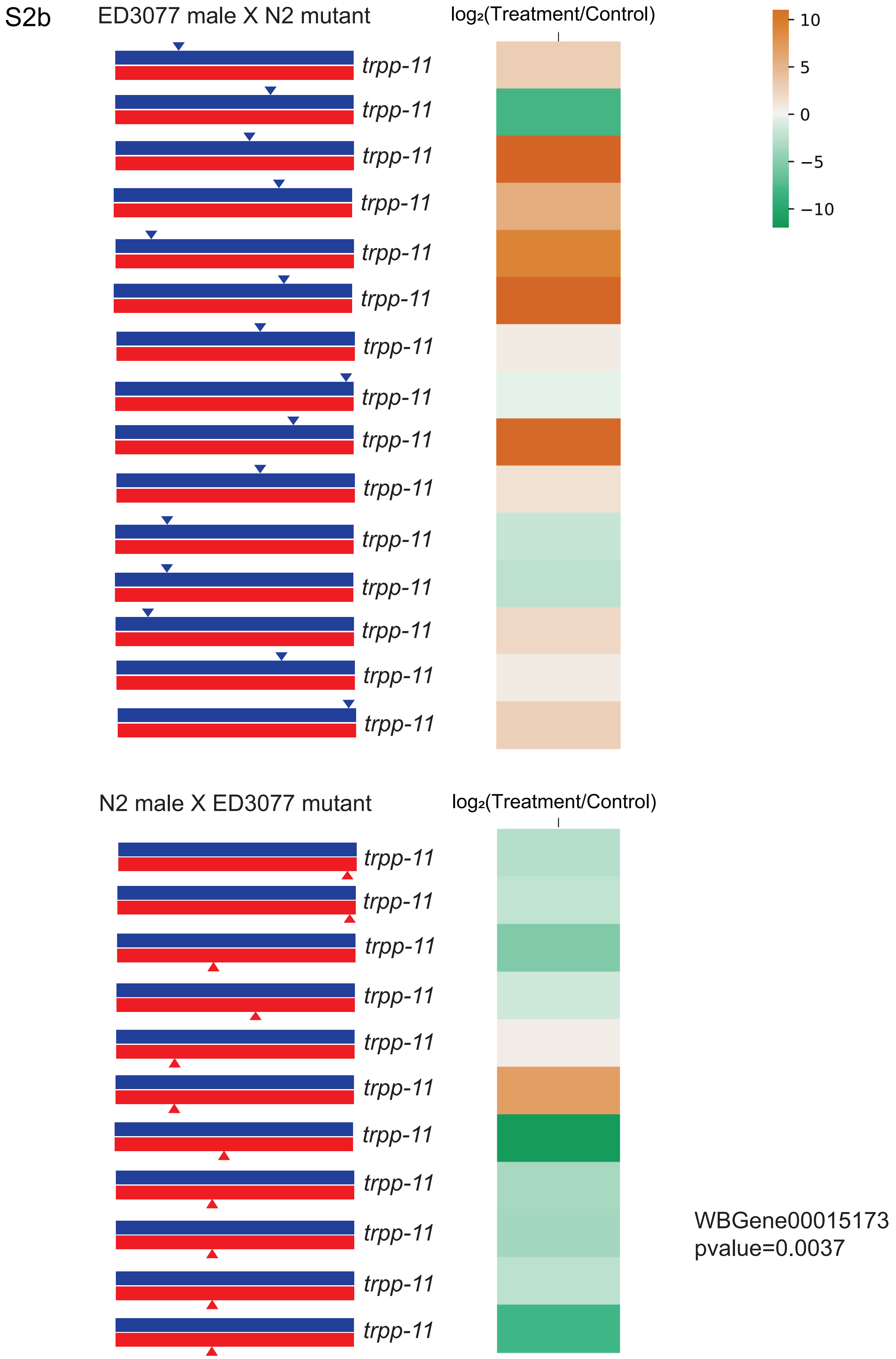

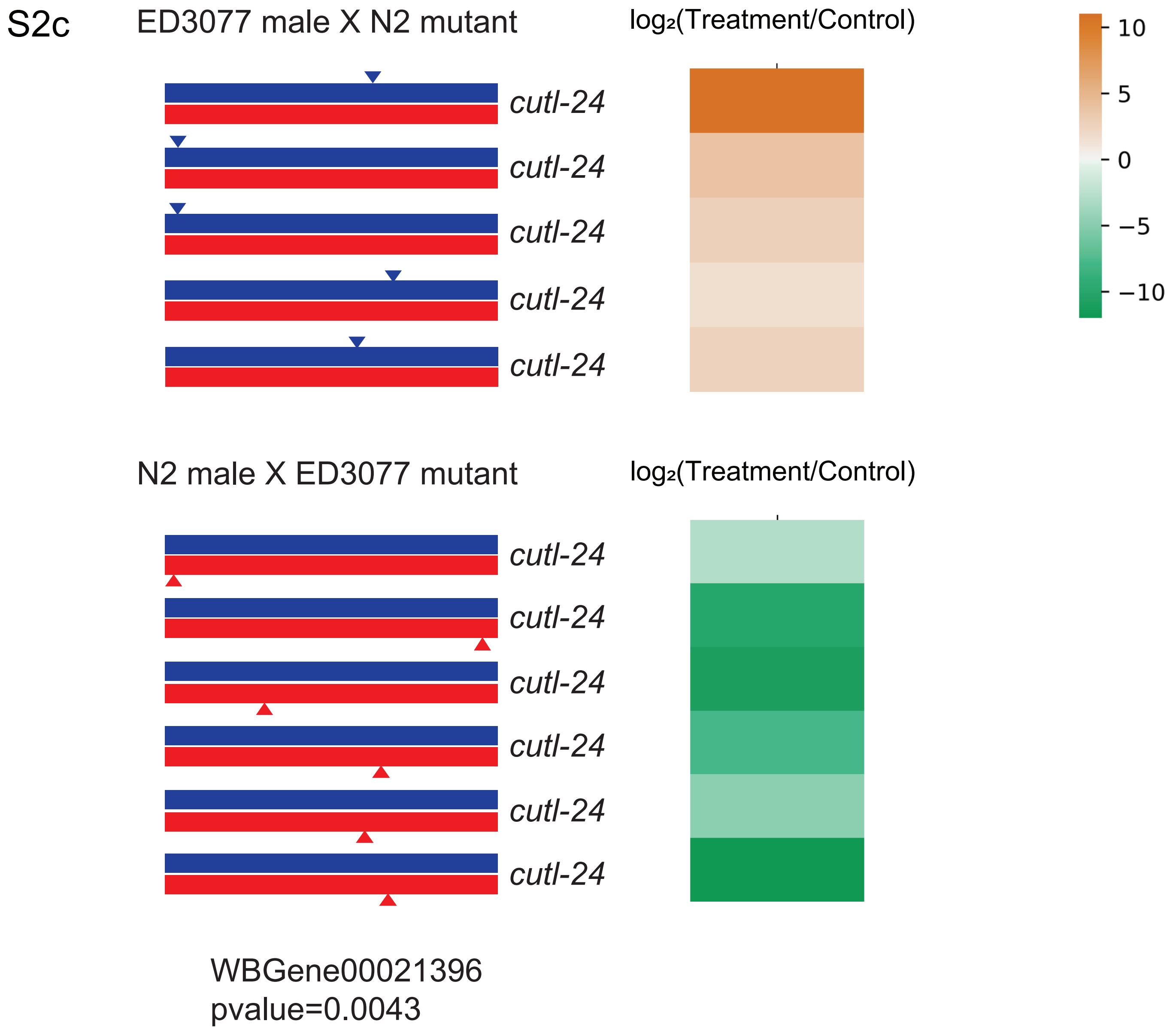

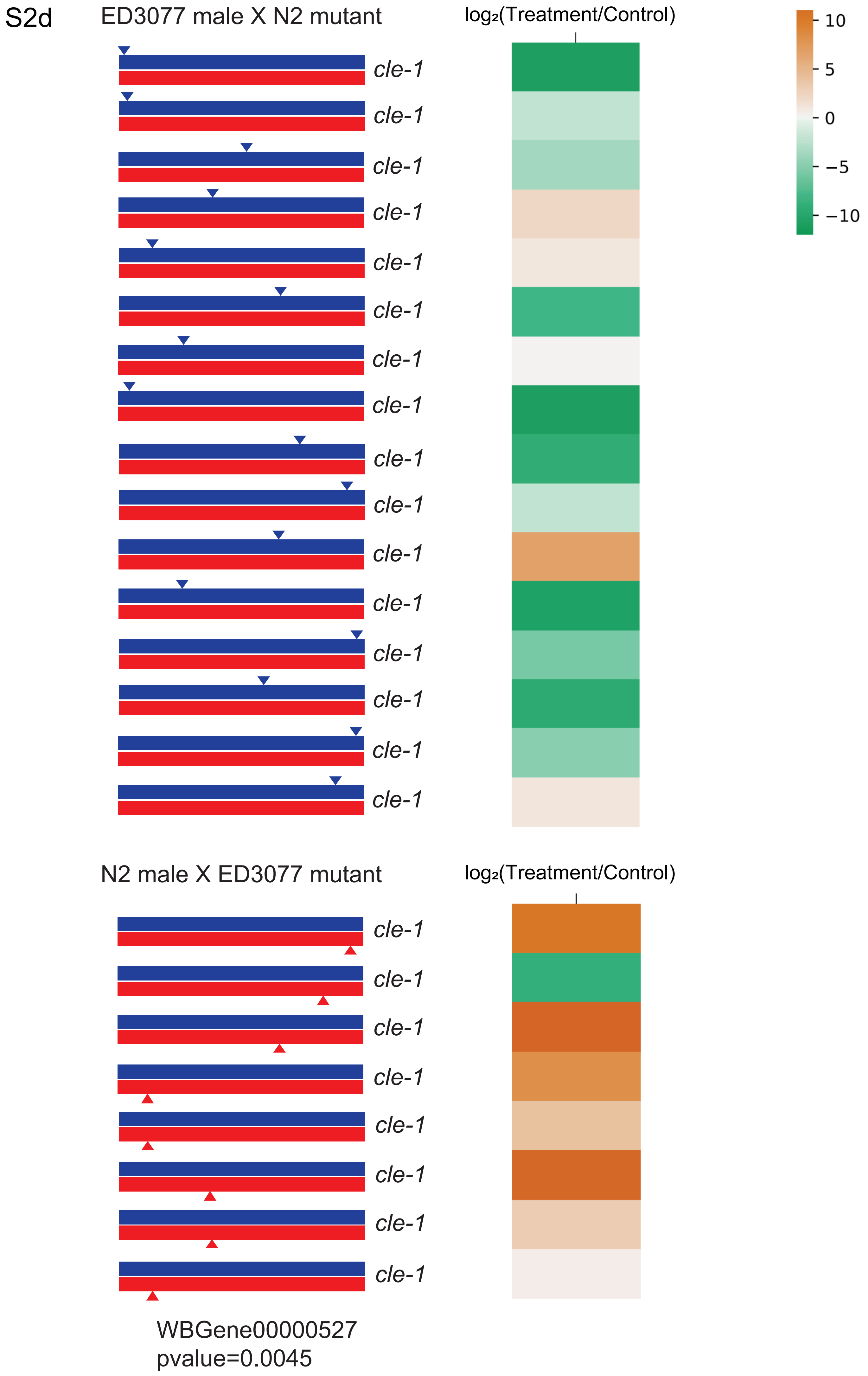

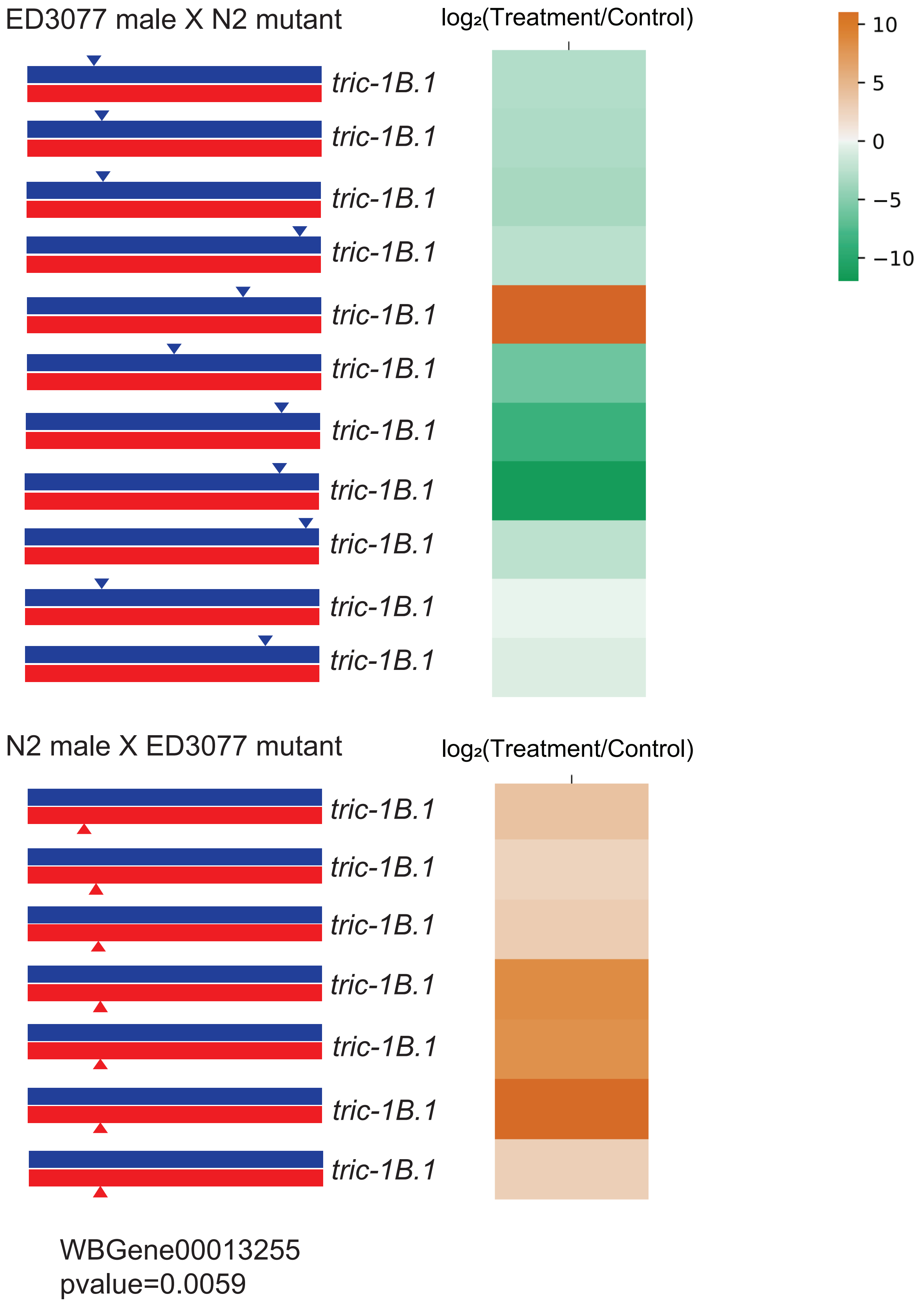

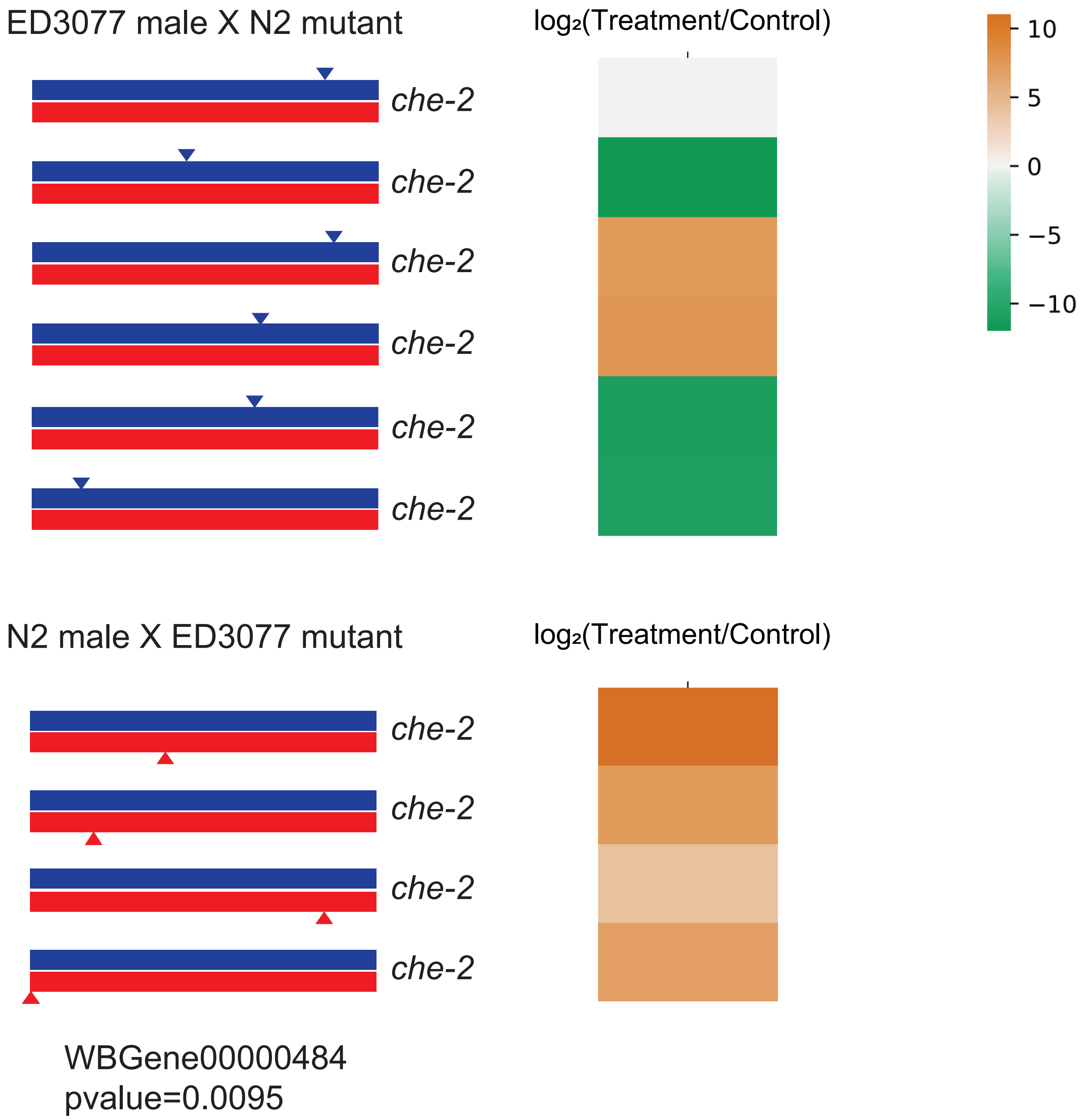

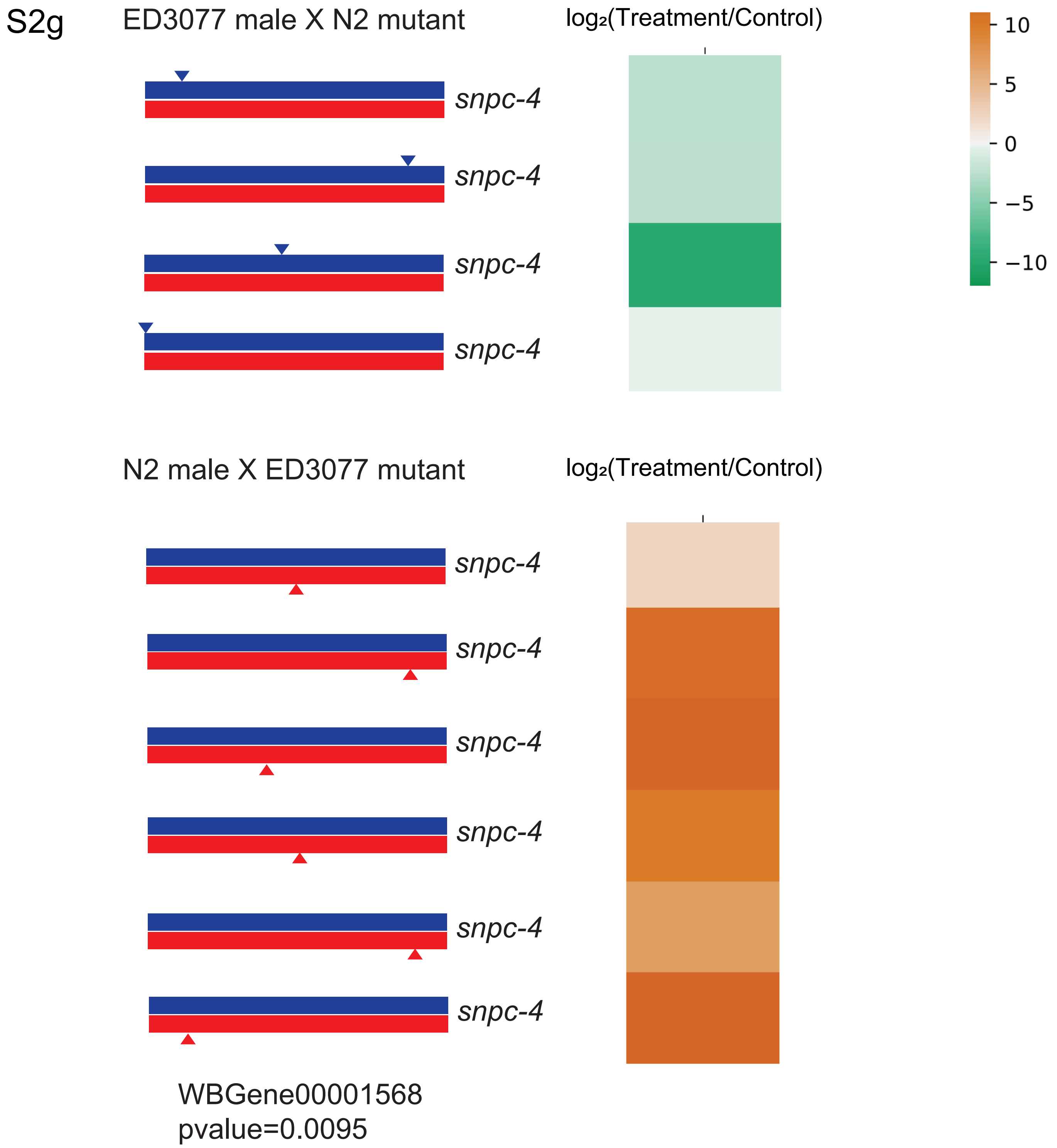

S2h

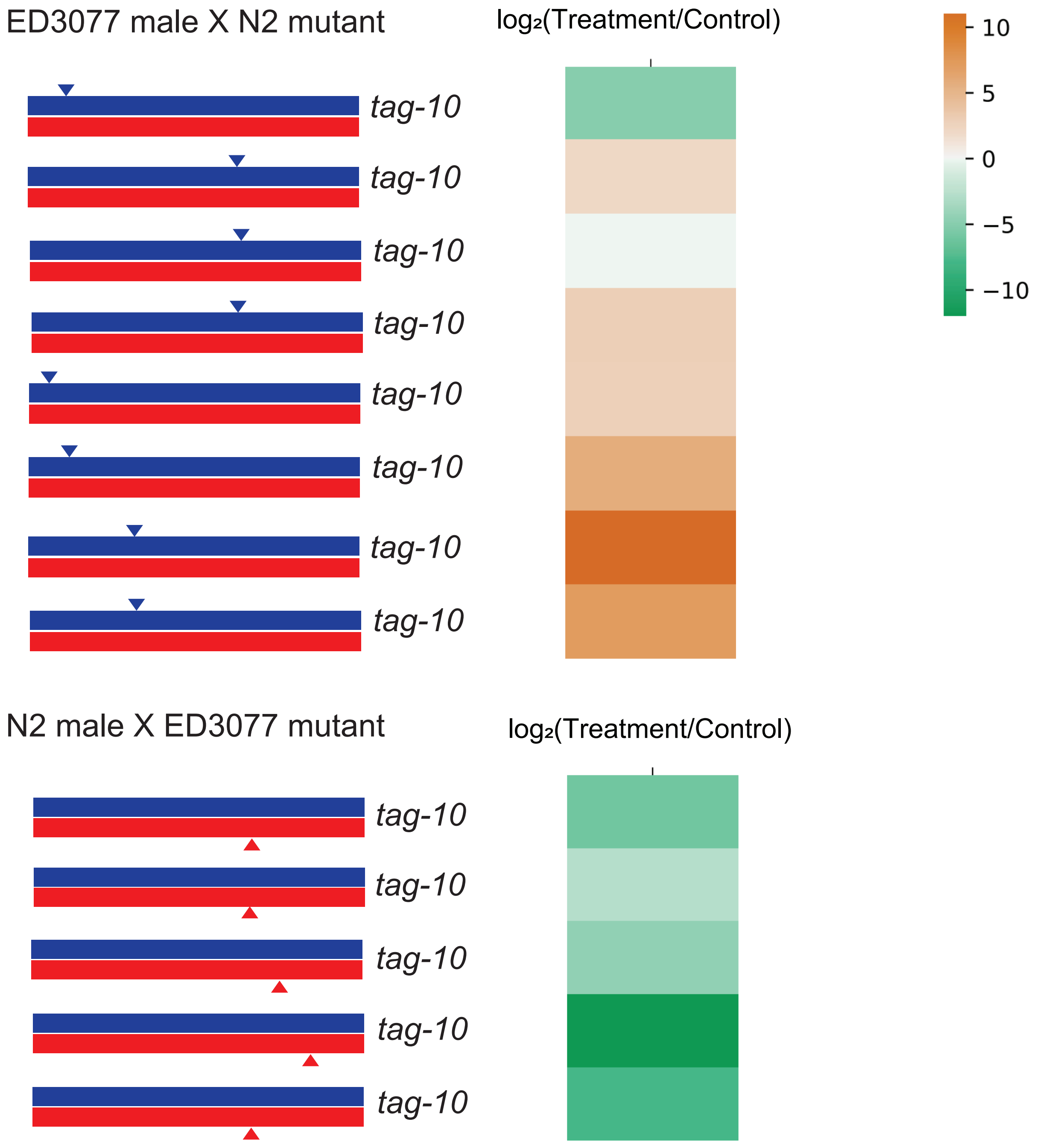


WBGene00006404
pvalue=0.011

S2i

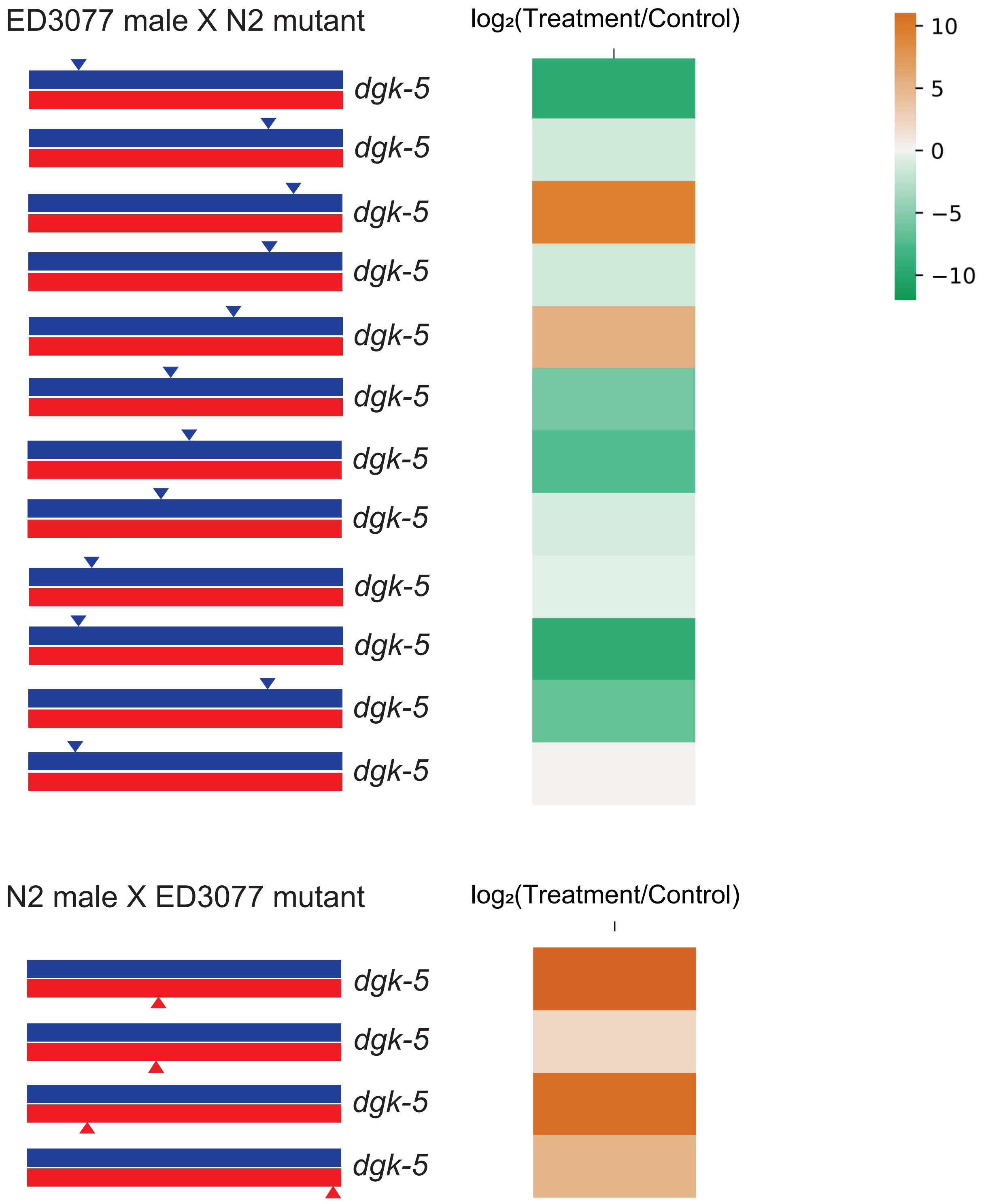


WBGene00019428
pvalue=0.013

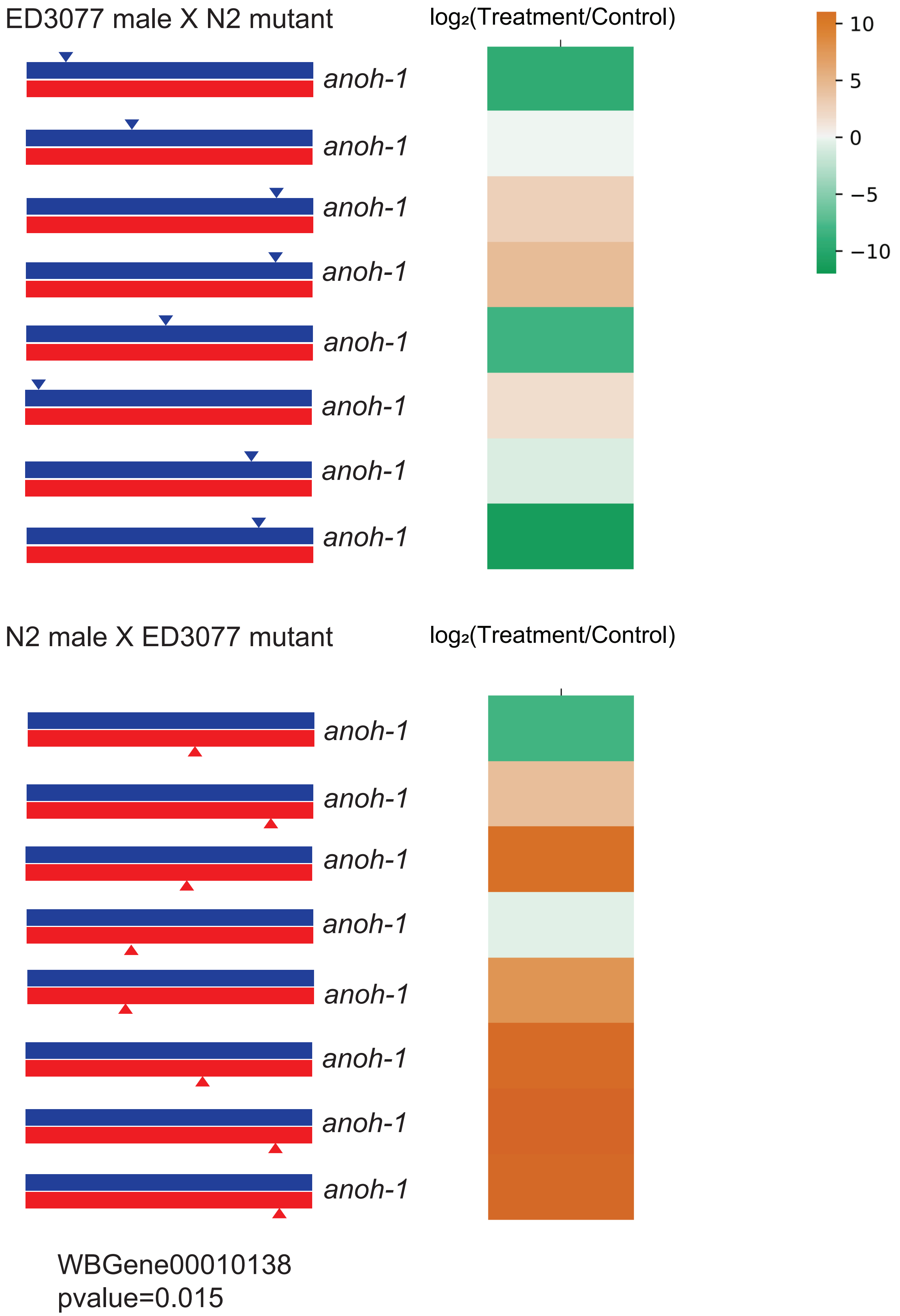
